## Supplementary Information for "Robust total X-ray scattering workflow to study correlated motion of proteins in crystals"

Steve P. Meisburger[1] David A. Case[2] Nozomi Ando[1*]

[1] Department of Chemistry and Chemical Biology, Cornell University, Ithaca, New York 14850, USA.

[2] Department of Chemistry and Chemical Biology, Rutgers University, Piscataway, New Jersey 08854, USA.

**Supplementary Table 1. Data collection parameters**

| Space group | Crystal | Sweep | Frames | Rotation (deg.) | Image file prefix |
|---|---|---|---|---|---|
| $P2_12_12_1$ | 1 | 1 | 1–500 | 0–50 | lys_rt_2_38 |
| | | 2 | 1–500 | 90–140 | lys_rt_2_39 |
| | | 3 | 1–500 | 180–230 | lys_rt_2_40 |
| | | 4 | 1–500 | 270–320 | lys_rt_2_41 |
| | 2 | 1 | 1–500 | 0–50 | lys_rt_5_51 |
| | | 2 | 1–500 | 90–140 | lys_rt_5_52 |
| | | 3 | 1–500 | 180–230 | lys_rt_5_53 |
| | | 4 | 1–500 | 270–320 | lys_rt_5_54 |
| $P3_32_12$ | 1 | 1 | 1–500 | 247–297 | lys_1_2 |
| | | 2 | 1–500 | 292–342 | lys_1_3 |
| | | 3 | 1–500 | 237–387 | lys_1_4 |
| | | 4 | 1–500 | 22–72 | lys_1_5 |
| | | 5 | 1–500 | 67–117 | lys_1_6 |
| | | 6 | 1–500 | 112–162 | lys_1_7 |
| | | 7 | 1–500 | 157–207 | lys_1_8 |
| | | 8 | 1–500 | 202–252 | lys_1_9 |

**Supplementary Table 2. Bragg data collection and structure refinement statistics**. *Values in parentheses refer to the highest resolution shell.

| | PDB ID: *8dz7* | PDB ID: *8dyz* |
|---|---|---|
| **Data Collection** | | |
| Space Group | P $2_1 2_1 2_1$ | P $4_3 2_1 2$ |
| Cell dimensions | | |
| $a$, $b$, $c$ (Å) | 30.49, 56.40, 73.85 | 79.63, 79.63, 38.30 |
| $\alpha$, $\beta$, $\gamma$ (°) | 90, 90, 90 | 90, 90, 90 |
| Resolution (Å) | 44.87–1.34 (1.36–1.34)* | 39.85–1.27 (1.29–1.27) |
| $R_{\text{merge}}$ | 0.039 (0.238) | 0.046 (0.380) |
| $R_{\text{pim}}$ | 0.011 (0.170) | 0.009 (0.184) |
| $I$ / $\sigma I$ | 36.0 (3.8) | 39.2 (3.7) |
| $CC_{1/2}$ | 0.999 (0.898) | 1.000 (0.878) |
| Completeness (%) | 96.9 (68.4) | 98.0 (78.1) |
| Multiplicity | 12.1 (2.2) | 23.8 (4.6) |
| **Refinement** | | |
| Resolution (Å) | 44.87–1.34 | 39.85–1.27 |
| Unique reflections: all/free | 28632 / 1417 | 32215 / 1659 |
| $R_{\text{work}}$ / $R_{\text{free}}$ | 0.118 / 0.136 | 0.115 / 0.133 |
| Number of non-H atoms | | |
| Protein | 1026 | 1030 |
| Ligand/ion | 2 | 4 |
| Water | 50 | 78 |
| Mean isotropic $B$-factors | | |
| Protein | 24.80 | 25.45 |
| Ligand/ion | 24.55 | 40.2 |
| Water | 34.32 | 35.76 |
| Model validation | | |
| Ramachandran outliers (%) | 0.00 | 0.00 |
| Ramachandran favored (%) | 99.21 | 99.21 |
| Rotamer outliers (%) | 0.00 | 0.00 |
| C-$\beta$ deviations | 0 | 0 |
| R.m.s. bond lengths (Å) | 0.0175 | 0.0160 |
| R.m.s. bond angles (°) | 1.97 | 1.85 |
| Clashscore | 0.00 | 0.00 |
| Overall score | 0.5 | 0.5 |

**Supplementary Table 3. Diffuse data processing and lattice disorder model statistics**

| | Triclinic | Orthorhombic | Tetragonal |
|---|---|---|---|
| | (P 1) | (P $2_12_12_1$) | (P $4_32_12$) |
| **Diffuse Map** | | | |
| Laue Group | *-1* | *mmm* | *4/mmm* |
| Reciprocal cell dimensions | | | |
| a*, b*, c* (Å$^{-1}$) | 0.0416, 0.0337, 0.0307 | 0.0328, 0.0177, 0.0135 | 0.0126, 0.0126, 0.0261 |
| $\alpha$, $\beta$, $\gamma$ (°) | 83.82, 70.60, 67.35 | 90, 90, 90 | 90, 90, 90 |
| Resolution (Å) | 25–1.25 | 20–1.28 | 33–1.16 |
| Voxel size (a*, b*, c*) | 1/13, 1/11, 1/11 | 1/13, 1/7, 1/5 | 1/5, 1/5, 1/11 |
| Merged observations ($\times 10^6$) | 43.9 | 13.2 | 9.9 |
| Completeness (%) | 97.8 | 92.0 | 89.0 |
| **DISCOBALL** | | | |
| 3D-ΔPDF | | | |
| Resolution cutoff (Å) | 1.25 | 1.62 | 1.62 |
| Voxel size (a, b, c) | 1/45, 1/54, 1/56 | 1/40, 1/70, 1/96 | 1/100, 1/100, 1/48 |
| Deconvolution | | | |
| Number of peaks in the asu | 787 | 84 | 36 |
| Radius cutoff (Å) | 4.0 | 4.0 | 4.0 |
| Resolution range (Å) | 5–1.67 | 5–1.67 | 5–1.67 |
| **GOODVIBES** | | | |
| Network model | | | |
| Supercell (a, b, c) | 13, 11, 11 | 13, 7, 5 | 5, 5, 11 |
| Rigid groups (per unit cell) | 1 | 4 | 8 |
| Contacts (per asu) | 310 | 232 | 202 |
| Unique Contacts | 155 | 116 | 102 |
| Reference halos | | | |
| Resolution range (Å) | 2.5–2.0 | 2.5–2.0 | 2.5–2.0 |
| Number of halos | 400 | 400 | 400 |
| Number of voxels | 621,858 | 166,773 | 101,265 |
| Simulation | | | |
| Resolution range (Å) | ∞–1.25 | ∞–1.62 | ∞–1.62 |
| Atomic displacement parameters | | | |
| Protein center of mass (Å) | -1.01, 14.46, 24.33 | -2.84, 12.51, -15.84 | -0.59, 20.68, 19.41 |
| Center of reaction (Å) | -1.85, 13.09, 25.77 | -3.15, 12.70, -12.80 | 2.33, 22.82, 21.15 |

| | Triclinic | Orthorhombic | Tetragonal |
|---|---|---|---|
| $T_{1,1}$, $T_{2,2}$, $T_{3,3}$ (Å$^2$) | 0.0362, 0.0450, 0.0350 | 0.1091, 0.0855, 0.1126 | 0.0942, 0.1138, 0.1278 |
| $T_{1,2}$, $T_{1,3}$, $T_{2,3}$ (Å$^2$) | -0.0003, 0.0040, 0.0022 | -0.0120, 0.0023, -0.0085 | 0.0041, 0.0124, -0.0100 |
| $L_{1,1}$, $L_{2,2}$, $L_{3,3}$ (deg.$^2$) | 0.5116, 0.5487, 0.5797 | 1.1164, 1.2407, 1.4753 | 1.7048, 1.3492, 1.1318 |
| $L_{1,2}$, $L_{1,3}$, $L_{2,3}$ (deg.$^2$) | -0.0196, -0.0566, -0.1044 | -0.1860, 0.2218, -0.0708 | -0.5496, 0.3593, -0.0113 |
| $S_{1,1}$, $S_{2,2}$, $S_{3,3}$ (deg. Å) | 0.0070, 0.0029, -0.0078 | -0.0426, 0.0255, 0.0129 | 0.0244, -0.0034, -0.0149 |
| $S_{1,2}$, $S_{1,3}$, $S_{2,3}$ (deg. Å) | -0.0061, 0.0034, -0.0004 | 0.0267, 0.0369, 0.0076 | -0.0184, 0.0015, 0.0383 |

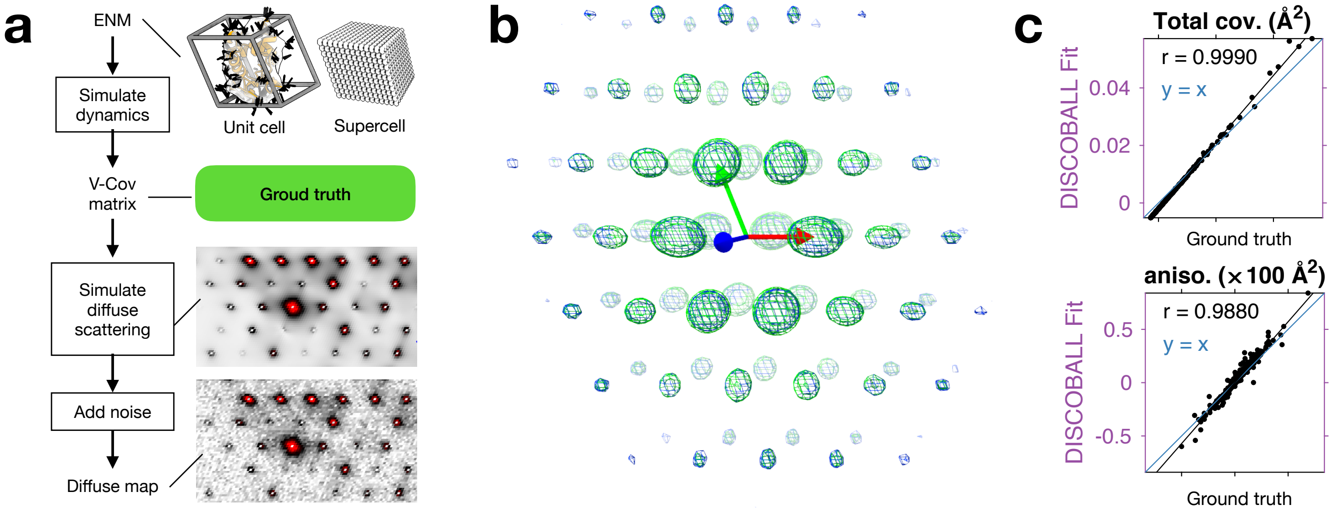

**Supplementary Figure 1. Validation of GOODVIBES models using DISCOBALL**. **a**, A synthetic diffuse scattering dataset of lysozyme in space group P1 was generated using a GOODVIBES ENM model with random spring constants. The ground truth joint-ADPs were calculated from the variance-covariance matrix (V-Cov). Uniform Gaussian random noise was added to the map, and DISCOBALL was applied to estimate the joint-ADPs. **b**, Overlay of the true (green) and DISCOBALL (blue) joint-ADPs represented using isosurface ellipsoids. **c**, Quantitation of DISCOBALL's ability to estimate joint-ADP ellipsoids. DISCOBALL recovers both the overall displacement covariance (top panel) and the anisotropic components (bottom panel) with high correlation ($r$, Pearson correlation coefficient).

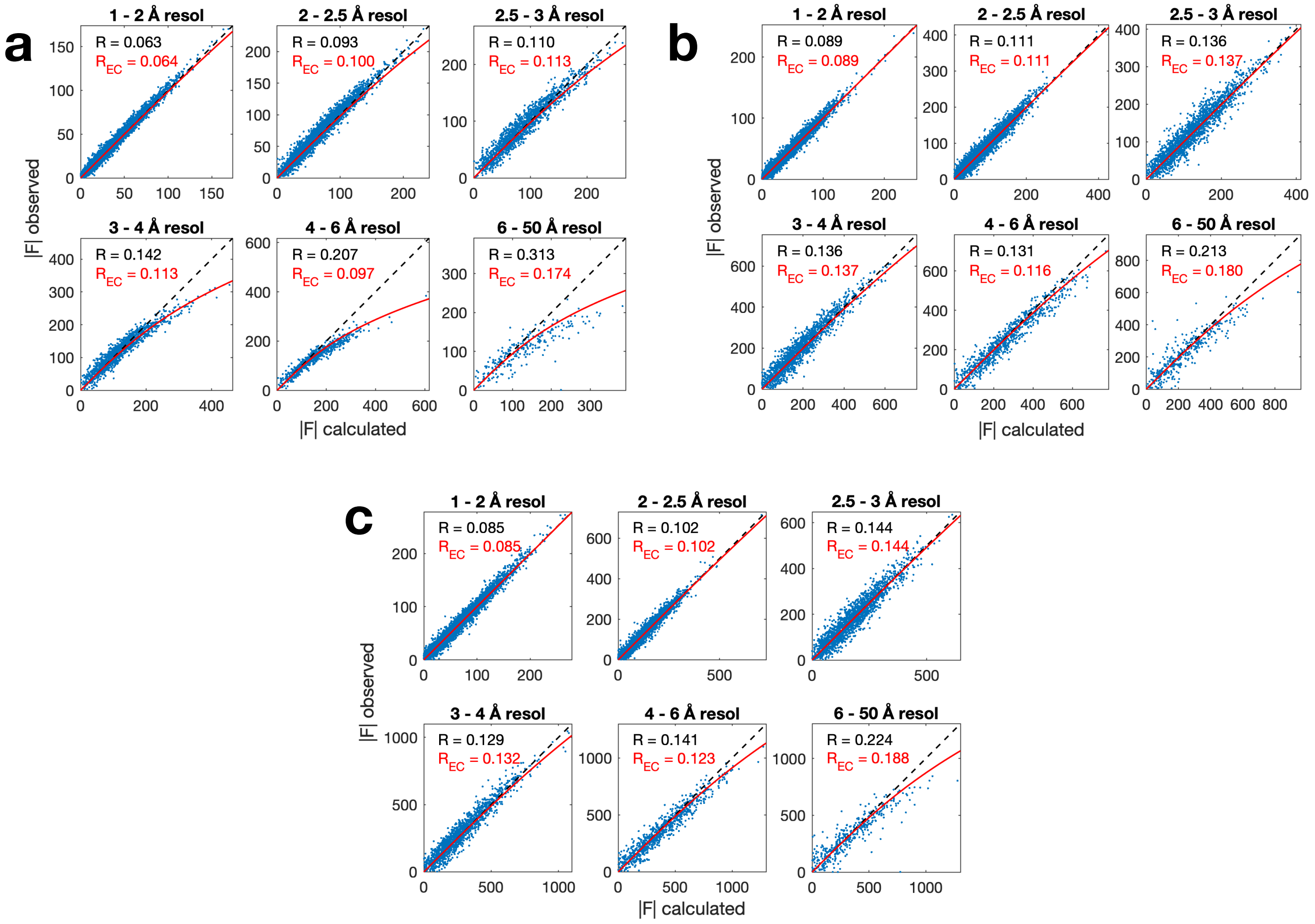

**Supplementary Figure 2. Apparent extinction of Bragg intensities from lysozyme polymorphs**. Accurate structure factor amplitudes are needed for GOODVIBES simulations of diffuse scattering. We therefore investigated the source of discrepancies between observed structure factors (Fobs) and those calculated from the refined structure (Fcalc). **a**, Scatter plots of Fobs vs. Fcalc for triclinic lysozyme grouped by resolution (indicated above each panel). Especially at low resolution (bottom row), Fobs is systematically lower than Fcalc in an intensity-dependent manner (points fall below the dashed line in each panel). This discrepancy leads to high R-factors at low resolution (insets). The discrepancy fits the functional form of the extinction correction (EC) used in small molecule crystallography[1] when its free parameter $x$ is optimized (red line, $x = 0.002$). After applying the EC, R-factors drop dramatically ($R_{EC}$, inset). **b**, Discrepancies between Fobs and Fcalc are also significant in orthorhombic lysozyme, but less so than in triclinic. Modest improvements in R-factor are seen when EC is applied ($x$=0.0001). **c**, Tetragonal lysozyme also shows slight discrepancies between Fobs and Fcalc at low resolution, and EC ($x$=0.00005) results in lower R-factors. Detector count-rate artifacts[2] might also explain the suppressed Bragg intensities, and cannot be ruled out without further experiments.

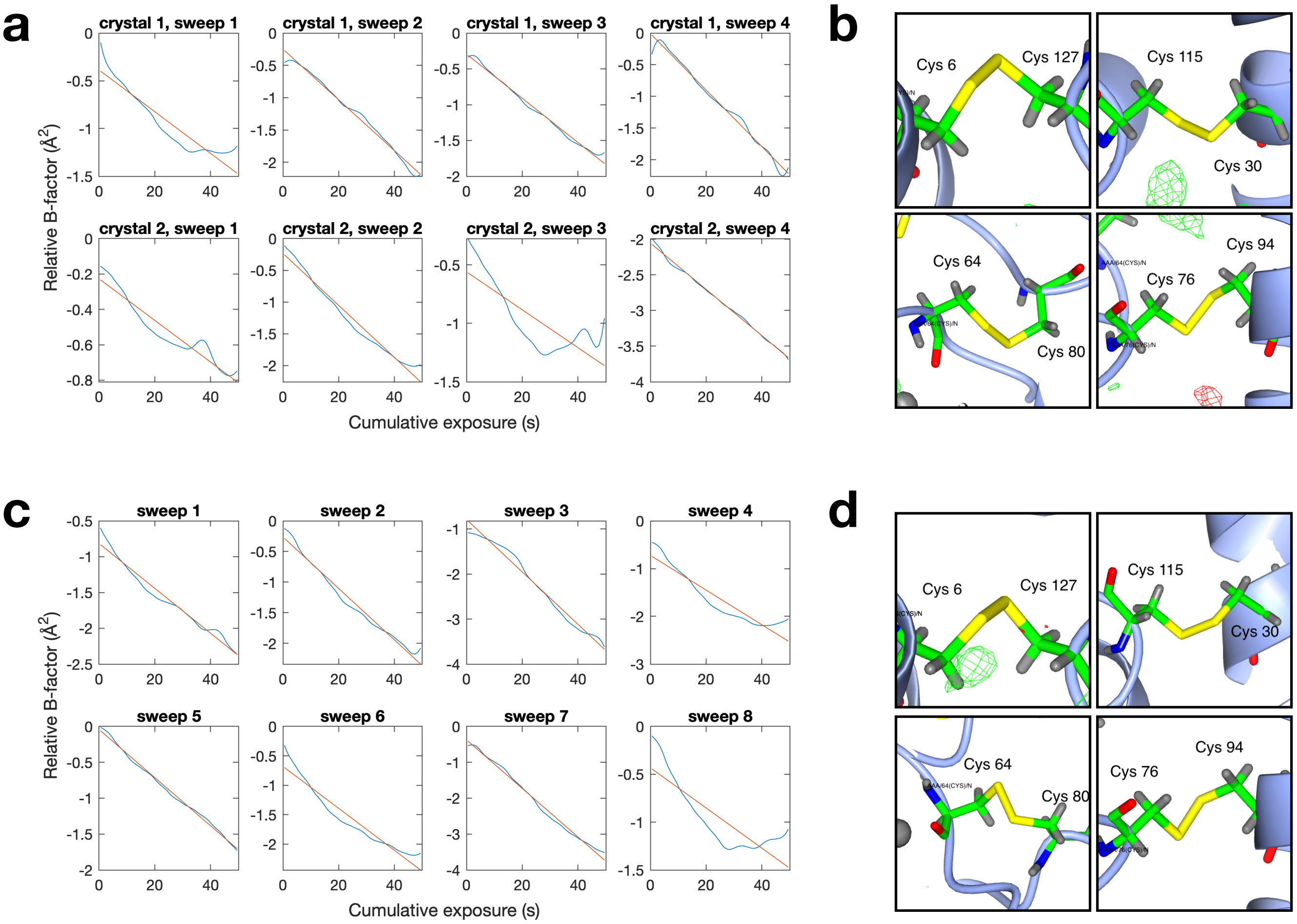

**Supplementary Figure 3. Monitoring global and site-specific radiation damage**. Global damage was assessed using B-factor correction applied during scaling, and site-specific damage was assayed using the difference electron density around disulfide bonds. **a**, B-factor decay profiles of orthorhombic lysozyme reported by *Aimless*[3] (blue lines) show a characteristic linear decay with X-ray exposure (red fits). **b**, Difference density (fo-fc) is not observed near the four disulfide bonds of orthorhombic lysozyme at the $3\sigma$ level (green and red mesh) signifying that all four disulfides remained intact. **c**, B-factor decays for the tetragonal dataset are similar to orthorhombic (panel a). **d**, Little difference density is observed near the disulfide bonds of tetragonal lysozyme, except for a small positive feature near Cys 6 may signify a minor population with reduced sulfhydryl groups (not modeled).

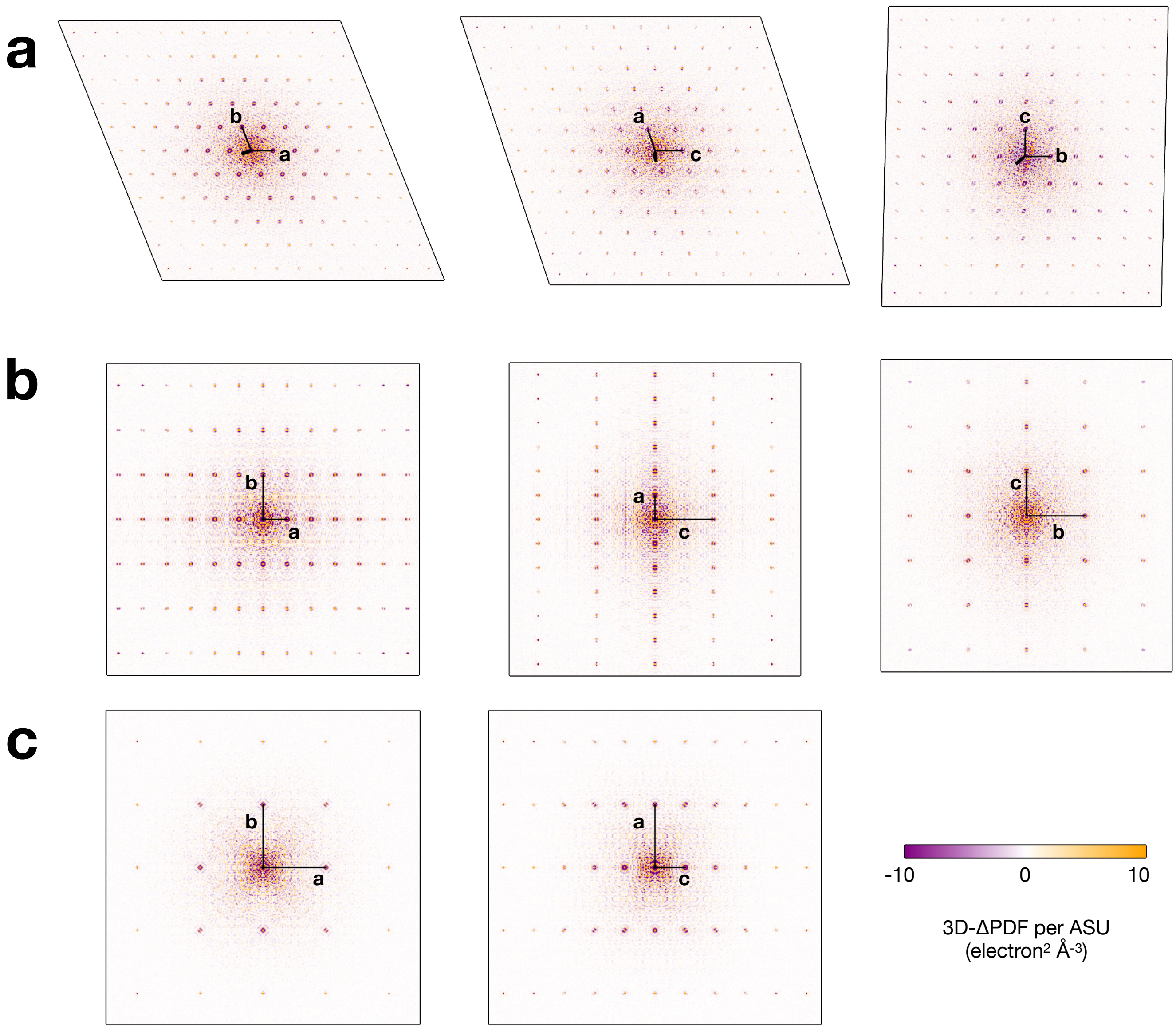

**Supplementary Figure 4. 3D-ΔPDFs of lysozyme polymorphs.** The 3D-ΔPDF (Fourier transform of the diffuse scattering) was calculated for three experimental datasets from lysozyme as described in Methods. Central sections through the 3D-ΔPDF for crystals in the triclinic (panel **a**), orthorhombic (panel **b**), and tetragonal (panel **c**) space groups show intense features near the origin and a series of sharp peaks that decay in intensity away from the origin. The sharp peaks occur at nodes of the direct lattice (multiples of the unit cell vectors **a**, **b**, and **c**). All maps are shown on the same scales of relative distance and intensity per asymmetric unit (ASU).

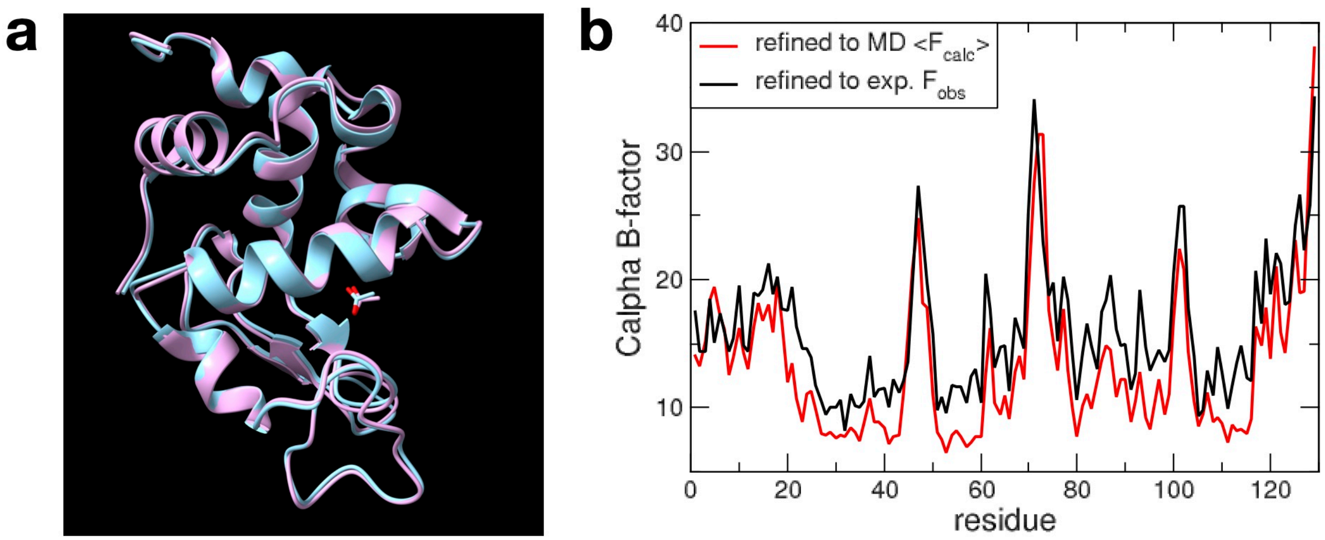

**Supplementary Figure 5. Comparison of MD simulation and experimental structures of lysozyme in tetragonal space group**. **a**, The deposited structure is cyan, refined structure using the MD average structure factors is pink. **b**, Comparison between the refined B-factors for C$\alpha$ atoms. The pattern is similar, although values refined against the MD data are somewhat lower in regions of secondary structure than are the values refined from experiment. Overall, these comparisons are very similar to those reported earlier for similar simulations of triclinic lysozyme[4].

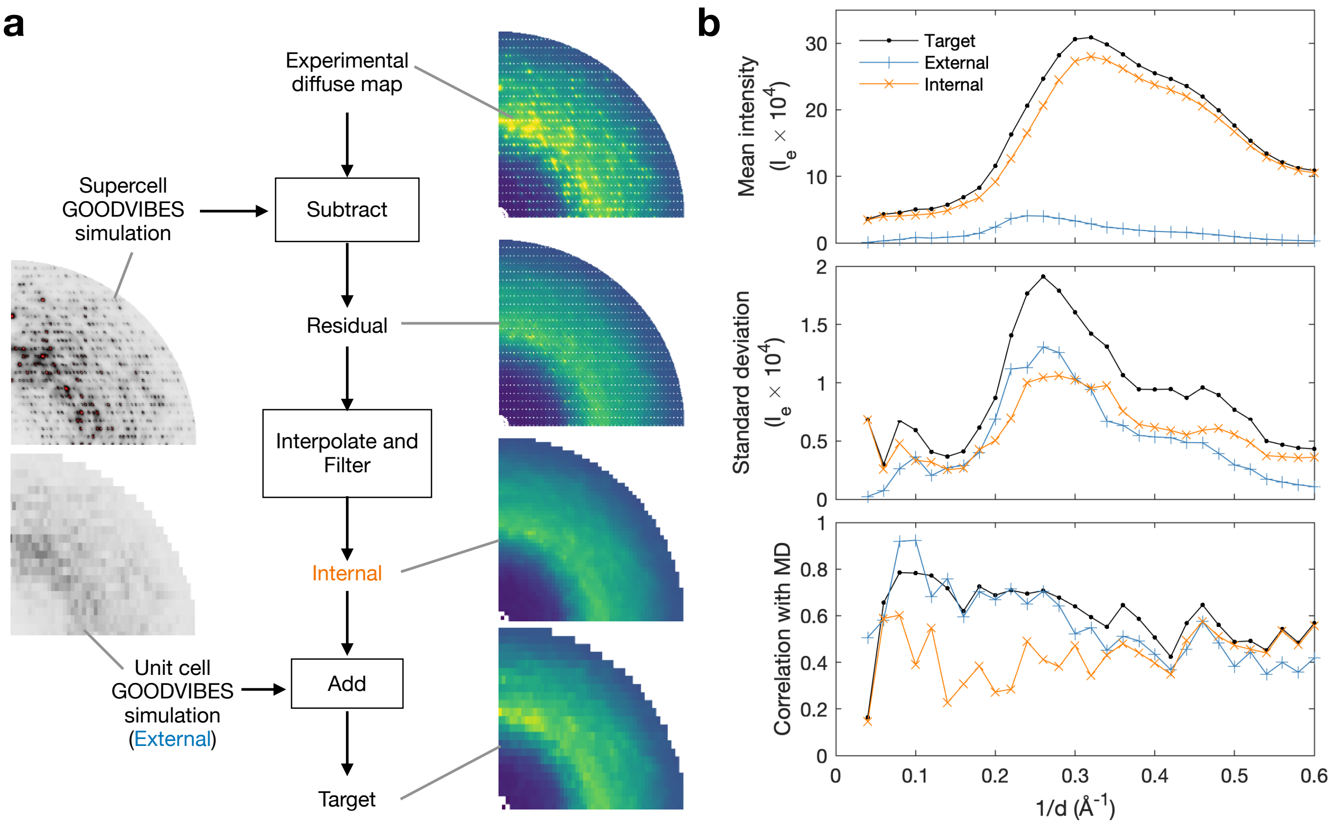

**Supplementary Figure 6. Components of the target diffuse map.** **a**, Procedure for producing target diffuse maps illustrated using tetragonal lysozyme from an experimental diffuse map (top right panel). First a GOODVIBES simulation of lattice dynamics (left panel) is subtracted, and then a filter is applied in order to interpolate this residual signal onto an appropriate grid and remove outliers. External motions compatible with the MD simulation are simulated using GOODVIBES. Finally, the target diffuse map is computed as the sum of internal and external maps. See Methods in the Main Text for details. **b**, Contribution of signals from internal (orange lines and symbols) and external (blue lines and symbols) motions to the target map (black lines and symbols) for tetragonal lysozyme. Internal motion makes the main contribution to the isotropic signal (top panel), and the internal and external have a comparable contribution to the scattering variations (middle panel). The correlation coefficient with MD is dominated by the external signal at low resolution but internal motion becomes important at high resolution (bottom panel).

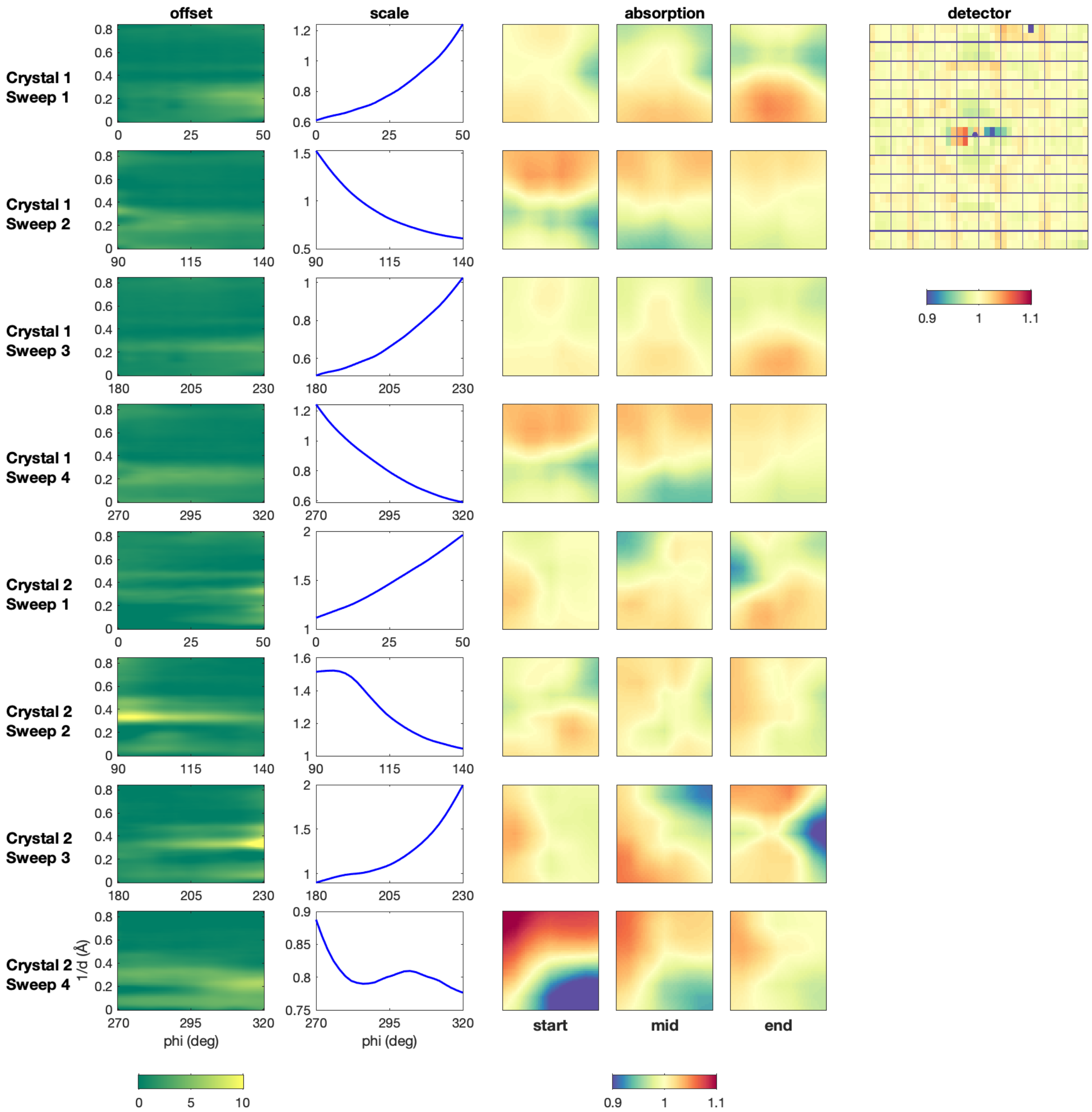

**Supplementary Figure 7. Refined scaling model to produce the diffuse map of orthorhombic lysozyme.** A scaling model was refined to correct for experimental artifacts in X-ray images. The model relates the merged intensity to the observed intensity as a function of spindle angle, resolution (d), and detector position. From left to right, the model parameters included: (1) offset correction vs. spindle angle and resolution, (2) overall scale factor vs. spindle angle, (3) scale factor for absorption correction vs. spindle angle and detector position, and (4) scale factor for detector efficiency correction vs. detector chip index. The parameters in the model were refined in order to minimize the least-squares error between observation and prediction. The best-fit parameters are illustrated for each partial dataset (top to bottom) (refer to Supplementary Table 1).

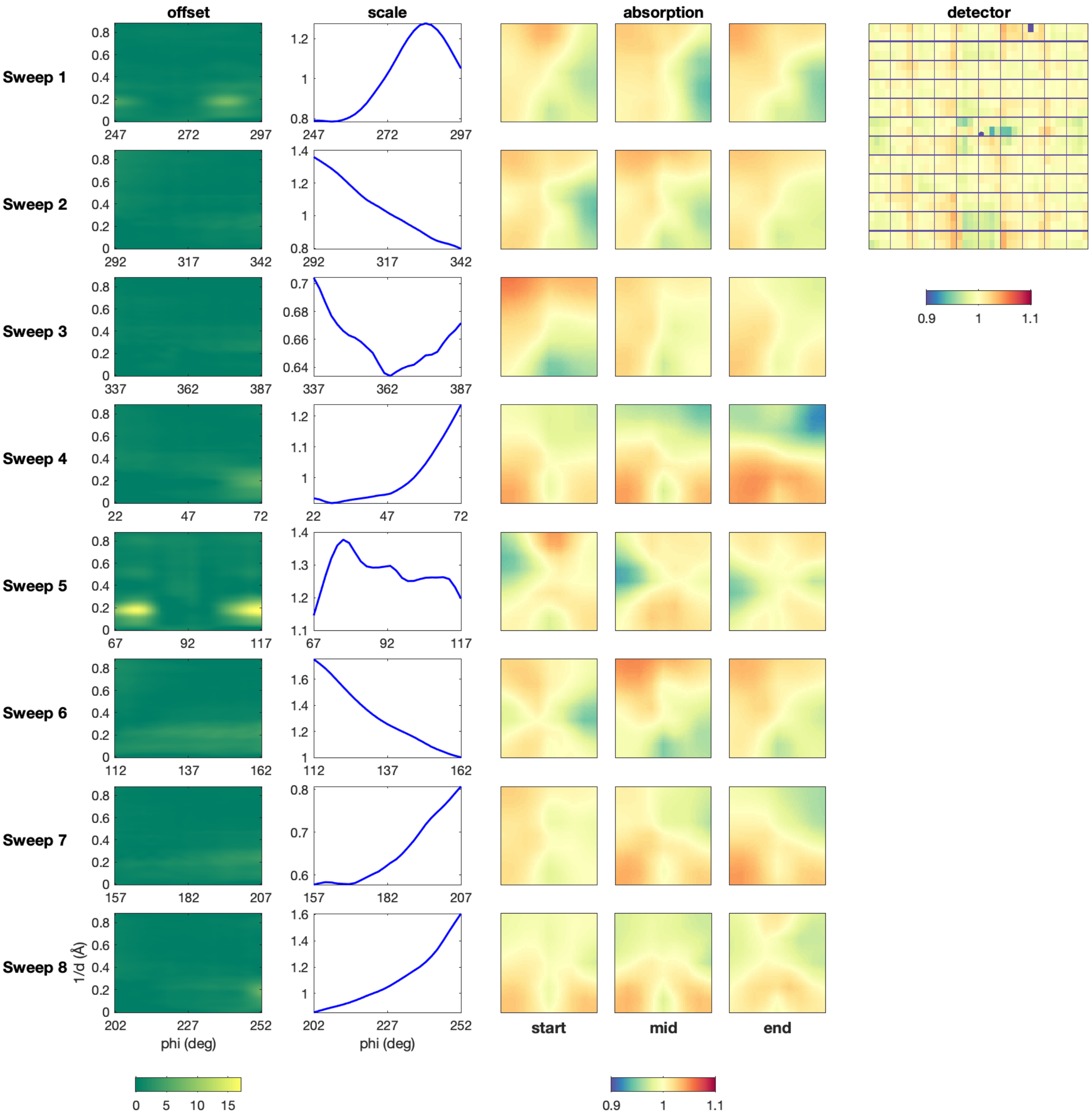

**Supplementary Figure 8. Refined scaling model to produce the diffuse map of tetragonal lysozyme.** For explanation, see Supplementary Fig. 7.

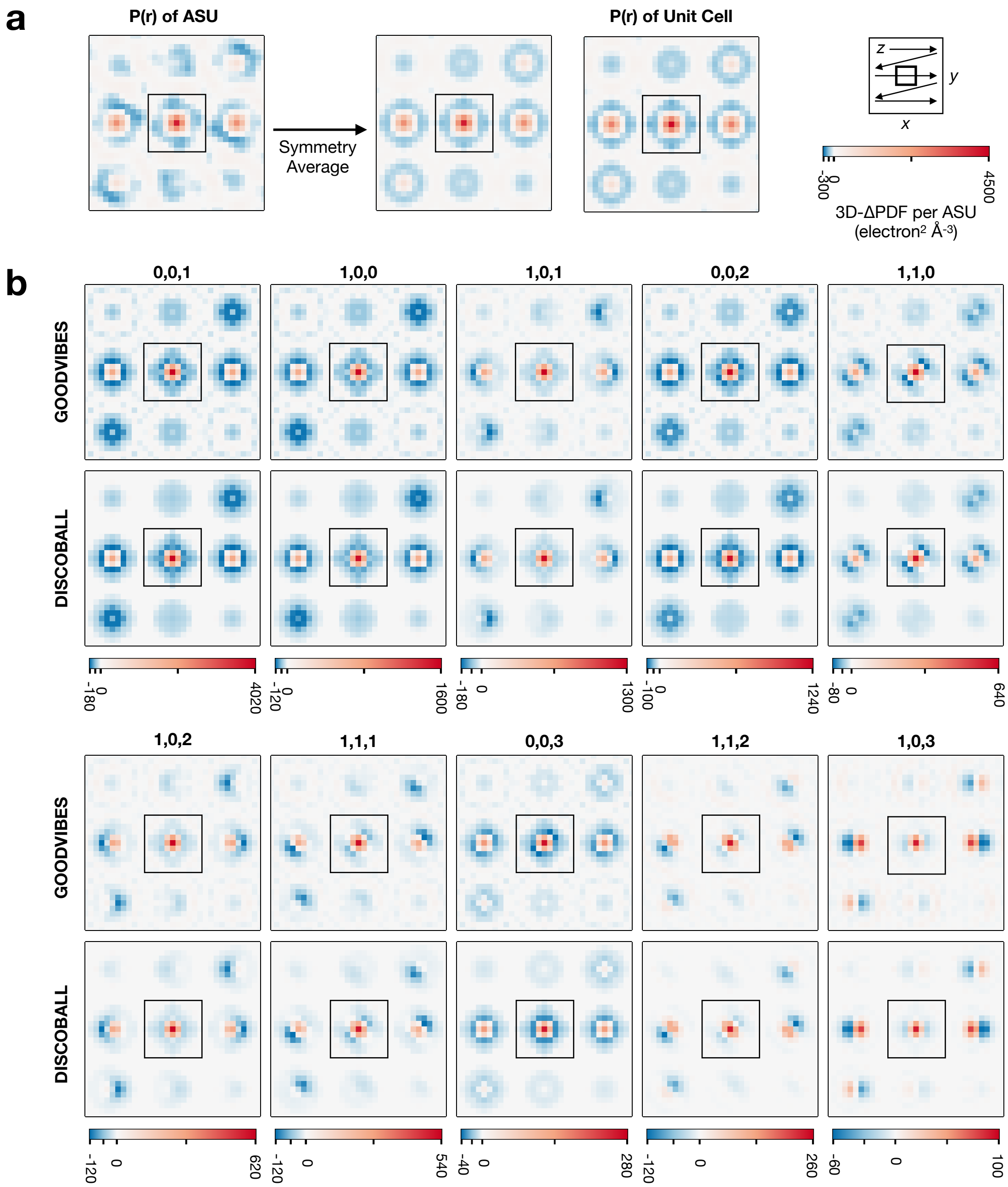

**Supplementary Figure 9. Test of approximations made in DISCOBALL model.** The DISCOBALL model for 3D-ΔPDF peaks is approximate for symmetric space groups (see Methods in the Main Text). Using tetragonal lysozyme as an example, we tested the effect of these approximations compared with an exact simulation. **a**, The central peak in the autocorrelation (P(r)) of the asymmetric unit (ASU) is approximately invariant to symmetry transformations (compare the red density in the left and central panels). Furthermore, the peak in the symmetry averaged asymmetric P(r) is nearly identical to that of the conventional Patterson function (P(r) of the unit cell, right panel), confirming that cross-terms between asymmetric units can be neglected. **b**, The 10 most intense 3D-ΔPDF peaks were simulated exactly using GOODVIBES and approximated using the DISCOBALL model with effective joint-ADPs computed from the GOODVIBES model (no deconvolution was performed). The similarity between the two confirms that approximations used in DISCOBALL have minimal effect on predicting the peak shape in this case.
